# Combination of novel and public RNA-seq datasets to generate an mRNA expression atlas for the domestic chicken

**DOI:** 10.1101/295535

**Authors:** Stephen J. Bush, Lucy Freem, Amanda J. MacCallum, Jenny O’Dell, Chunlei Wu, Cyrus Afrasiabi, Androniki Psifidi, Mark P. Stevens, Jacqueline Smith, Kim M. Summers, David A. Hume

## Abstract

**Background:** The domestic chicken (*Gallus gallus*) is widely used as a model in developmental biology and is also an important livestock species. We describe a novel approach to data integration to generate an mRNA expression atlas for the chicken spanning major tissue types and developmental stages, using a diverse range of publicly-archived RNA-seq datasets and new data derived from immune cells and tissues.

**Results:** Randomly down-sampling RNA-seq datasets to a common depth and quantifying expression against a reference transcriptome using the mRNA quantitation tool Kallisto ensured that disparate datasets explored comparable transcriptomic space. The network analysis tool Miru was used to extract clusters of co-expressed genes from the resulting expression atlas, many of which were tissue or cell-type restricted, contained transcription factors that have previously been implicated in their regulation, or were otherwise associated with biological processes, such as the cell cycle. The atlas provides a resource for the functional annotation of genes that currently have only a locus ID. We cross-referenced the RNA-seq atlas to a publicly available embryonic Cap Analysis of Gene Expression (CAGE) dataset to infer the developmental time course of organ systems, and to identify a signature of the expansion of tissue macrophage populations during development.

**Conclusion:** Expression profiles obtained from public RNA-seq datasets – despite being generated by different laboratories using different methodologies – can be made comparable to each other. This meta-analytic approach to RNA-seq can be extended with new datasets from novel tissues, and is applicable to any species.

## INTRODUCTION

Aggregation and meta-analysis of multiple large gene expression datasets based upon common microarray platforms is relatively commonplace in many species (e.g. [1–3]). Although RNA-seq is rapidly supplanting microarrays for gene expression profiling, it is not yet clear whether data from multiple different labs can be analysed together in an informative manner. Confounding variables reflect the many technical – and bias-prone – aspects of library preparation and sequencing (see reviews [4, 5]), with RNA-seq datasets often differing in read length [6], depth of coverage [7], strand specificity [8], RNA extraction and library selection methods [9], sequencing platform [10, 11] and the choice to sequence single- or paired-end reads [12]. For a given dataset, these variables can together affect both the number and type of genes detectable and the accuracy of their expression level estimates. Expression quantification is also affected by sample quality [13] and storage method [14], irrespective of sequencing technique: RNA degrades with lengthier post-mortem intervals [15] (the extent of which is tissue-dependent [16]) with degradation resulting in inaccurate quantification, particularly for shorter transcripts [17]. Sequencing composite biological structures (those with internal structures that have distinct functions), whether intentionally or inadvertently, can mask the signal of structure-specific differential expression [18]. Despite these variables, meta-analysis combining mammalian gene expression datasets [19–21] suggests that RNA-seq datasets are generally robust to inter-study variation, with the expression profiles of homologous tissues clustering more closely with each other than with different samples from the same study or species [22]. Expression atlases are valuable resources for functional genomics. Groups of transcripts – members of which will have similar expression profiles – can be associated with a shared function, such as a particular pathway or biological process. This principle is known as ‘guilt by association’ [23] and has previously been used to annotate genes of unknown function in human [2, 24, 25], pig [26], sheep [27] and mouse [28, 29] datasets. Co-expression information is also informative in genome-wide association studies (GWAS) of complex traits and disease susceptibility. The simple principle, that genes involved in the same trait or phenotype tend to be expressed in the same cell type or tissue, or otherwise participate in the same pathway, has been confirmed in multiple datasets [28, 30]. Because of the ease of access *in ovo*, the chicken (*Gallus gallus*) embryo has been widely used as a model system in cell and developmental biology, constrained only by methods for genomic manipulation *in situ*, or in the germ line. These constraints were largely overcome through the sequencing of the genome, and technological developments such as *in vivo* electroporation, more than 15 years ago [31, 32]. More recent innovations including the generation of reporter transgenes [33] and genome editing via primordial germ cells [34–36] have transformed the utility of the chicken as a model organism. However, the current genome build still has many unannotated or minimally annotated genes about which very little is known [28]. Of the 18,347 protein-coding genes in version GalGal5 of the chicken genome in Ensembl89, 7275 (40%) have only been assigned an Ensembl placeholder ID.

The domestic chicken is also a major source of animal protein worldwide, with different lines heavily selected for optimal production traits such as increased egg production or rapid weight gain. The molecular basis for these traits is increasingly being associated with genomic loci through genome-wide association studies based upon high density SNP platforms [37]. Both the application of the chick as a model organism, and for candidate gene analysis in genomic intervals associated with trait variation, would be expedited by improvements in functional genome annotation. In particular, it would be useful to identify the sets of protein-coding genes that share transcriptional regulation between the chick and the mouse, the most widely-studied mammalian model organism. For this purpose, we aimed to generate a comprehensive atlas of mRNA expression for the chicken.

With the removal of antibiotics from the food chain and threats from emerging diseases, there is also interest in the selection of birds with increased disease resistance and/or resilience [38]. To support this activity, we were particularly interested in identifying and annotating genes expressed specifically at high levels in cells of the innate immune system. Such gene sets have been identified in previous studies of human [2, 24, 25], pig [26], sheep [27] and mouse [28].

The current version of the chicken assembly was largely derived from high-throughput (i.e. comparatively cheap but imprecise) short read sequencing and primarily contains protein-coding gene models. The recent use of long-read – PacBio SMRT Iso-Seq – data has demonstrated that the transcriptomic complexity of chickens is comparable to humans, with many additional lncRNA models (among others) scheduled for inclusion in future Ensembl annotations [39].

To identify the set of genes expressed in innate immune cells in both unchallenged and activated conditions, we generated pure cultures of bone marrow-derived macrophages (BMDMs) grown in the presence of recombinant chicken macrophage colony-stimulating factor (CSF1), and stimulated them with the archetypal microbial agonist, lipopolysaccharide (LPS) [40]. To complement the data generated from macrophages *in vitro*, we also obtained RNA-seq libraries from the caecal tonsils of birds infected with *Campylobacter*, as well as from previous studies of macrophage, dendritic cell and heterophil populations. A global expression atlas for the chicken transcriptome was created by combining our immune-related data with 20 publicly archived RNA-seq datasets. Some were collated by the Avian RNA-seq Consortium [41], while others are drawn from a diverse range of existing publications, including studies that characterised the genetic basis of retinogenesis [42], the genetic determinants of meat tenderness [43], the morphological diversity of skin appendages [44], visceral fat metabolism [45], the transition between laying and brooding phases [46], the effect of heat stress upon pituitary development [47] and spleen function [48], the pathways involved in avian influenza resistance [49], the role of lncRNAs in the development of muscle [50], liver and adipose [51], and the transcriptional landscape of mRNA editing [52]. In total, 279 RNA-seq libraries were obtained, representing 48 distinct tissue and cell types at developmental stages spanning early embryonic (5 days) to mature adult (70 weeks post-hatching). In addition, we accessed a recently published transcriptional analysis of chick development generated by Cap Analysis of Gene Expression (CAGE) [53], a technique which can be used to quantify gene expression based on the transcript start site [54]. We show that the ‘guilt by association’ approach to functional annotation is viable even when combining disparate RNA-seq datasets, and utilise the meta-dataset to identify macrophage-specific and other informative co-expression clusters, providing a resource for genetic and genomic study of avian trait variation.

## RESULTS

### Selecting samples for inclusion in an RNA-seq meta-dataset

Many chicken RNA-seq datasets are available in public repositories, as detailed in [41]. Robust co-expression clustering of any two genes depends upon sampling tissues and cells in which both vary across the widest possible range. To maximise the co-expression signal, we chose datasets to represent the greatest possible diversity of tissues and organ systems. Not all studies contain links to a publicly archived dataset, such as a study of induced ochratoxicosis in the kidney cortex [55] and two studies of the bursa of Fabricius [56, 57]. Samples containing less than 10 million reads were not used, such as those from a study of the follicular transcriptome throughout the ovulation cycle [58].

Datasets used are detailed in Table S1, and have few commonalities: they were sequenced using a variety of Illumina instruments (HiSeq 2000/2500/3000/4000, Genome Analyzer II/IIx, NextSeq 500 and HiScanSQ), and include single- and paired-end, strand-specific and non-specific, polyA-selected (mRNA-seq) and rRNA-depleted (total RNA-seq) libraries at different read lengths and depths. For 12 tissues, independently sequenced RNA-seq datasets for the same tissue (Table S2) also allow for internal tests of the validity of aggregating the data. Throughout this text studies are referred to by their NCBI BioProject ID.

### Quantifying expression by iteratively revising a reference transcriptome

Expression was quantified – as transcripts per million (TPM) – using an RNA-seq processing pipeline [59] which iteratively runs the quantification tool Kallisto [60] with each iteration using an incrementally revised transcriptome. Kallisto requires that the user provide a set of transcripts, which are decomposed into k-mers. The expression of each transcript is quantified by matching this set of k-mers to the k-mers of the reads. For the first iteration of Kallisto, a non-redundant transcriptome (57,234 transcripts, representing 17,680 Ensembl protein-coding genes) was obtained by combining Ensembl transcript models with NCBI mRNA RefSeqs (see Materials and Methods).

The output was first parsed for library quality. The reverse cumulative distribution of TPM per gene was plotted on a log-log scale (Figure 1). The distributions generally approximate a power-law with an exponent of approximately -1 (Table S3), consistent with Zipf’s law (that the probability of an observation is inversely proportional to its rank) [61, 62]. Four samples with exponents < -0.8 or > - 1.2, i.e. deviating > 20% from the optimal value of -1 – were excluded from further analysis (i.e. the next iteration of Kallisto) (Table S3). Using only data from the useable samples, we created a revised reference transcriptome. During the first iteration of Kallisto, 55,027 of 57,234 transcripts (96%) were detectably expressed (average TPM > 1 in at least one tissue, where the average is the median TPM across all replicates, per BioProject, of that tissue), representing 17,313 Ensembl protein-coding genes (Table S4). After excluding 2207 transcripts with TPM < 1 in all tissues (Table S5) and those detectable only in the 4 excluded samples (n = 57), a revised transcriptome was generated containing 54,970 transcripts. For the second iteration of Kallisto, expression was re-quantified using this revised transcriptome, creating a final set of gene-level TPM estimates. The overall meta-dataset provides gene-level expression for 23,864 gene models (both Ensembl and NCBI) as median TPM across all replicates, per BioProject, per tissue (Table S6). Of these gene models, 43% (10,090) were unannotated, having only either an Ensembl placeholder ID or an NCBI locus ID.

**Figure 1.**
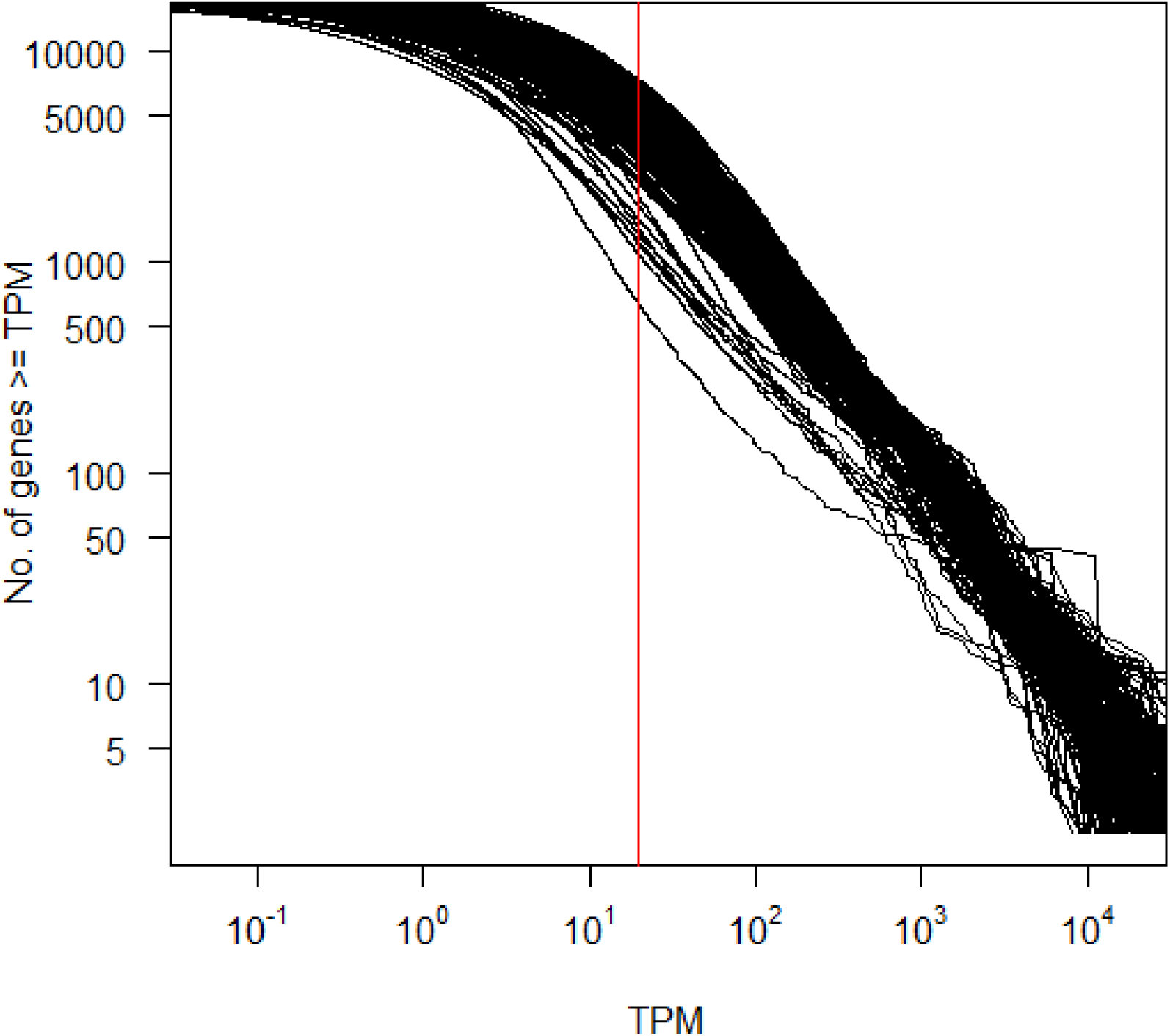
Reverse cumulative distribution of the number of genes that have at least a given TPM. Both axes are logarithmic. Each line represents data from an individual SRA sample ID, quantified using the first iteration Kallisto transcriptome (i.e. a non-redundant set of Ensembl protein-coding CDS plus trimmed RefSeq mRNAs). Samples are not otherwise distinguished as in general, most relationships approximate the same power-law: a minority of genes account for the majority of reads. These relationships are piecewise linear because the capture of lowly expressed genes is noisy, an artefact of random transcriptome sampling. The vertical red line denotes TPM = 5. At higher values of TPM, the majority of samples have a log-linear relationship. Those that do not are erroneous, and are excluded from subsequent analysis. Exponents of each sample’s log-log plot are given in Table S3.

### Randomly down-sampling RNA-seq datasets does not quantitatively alter their expression profiles

Higher resolution expression profiles are dependent upon higher sequencing depths [63] with diminishing returns – after approximately 10 million reads – on the power to detect genes differentially expressed between conditions [64]. For the purpose of functional annotation, it is more important to minimise variation between samples than to comprehensively capture transcripts. Accordingly, all datasets were randomly down-sampled to exactly 10 million reads before quantification.

To ensure the resulting co-expression signals are reproducible, it is necessary to establish that there are no significant differences in expression profiles introduced by sampling. For instance, the LPS-stimulated BMDM datasets were sequenced at depths of 37.5 to 52.6 million reads, such that when down-sampling, the BMDM expression profile as quantified for the meta-dataset was obtained using approximately one fifth to one quarter of the original reads (Table S7). To validate the approach, we randomly down-sampled each BMDM dataset to 10 million reads 100 times, using seqtk (https://github.com/lh3/seqtk, downloaded 29^th^ November 2016) seeded with a random integer between 0 and 10,000 (Dataset S1). After performing an all-against-all correlation of the 100 sets of data, the average Spearman’s *rho* was > 0.96 (Table S8), with the absolute difference, per gene, between maximum and minimum expression level averaging approximately 8 TPM (Figure 2 and Table S9). 70-75% of the genes detectably expressed (TPM > 1) in at least one of the 100 random samples were detected in all 100 samples (Table S8). Conversely, <5% of the genes were detectable in <5% of the samples (Table S8). The detection of these genes was stochastic, as they were expressed at very low levels – on average, 1.3 TPM (Table S8).

**Figure 2.**
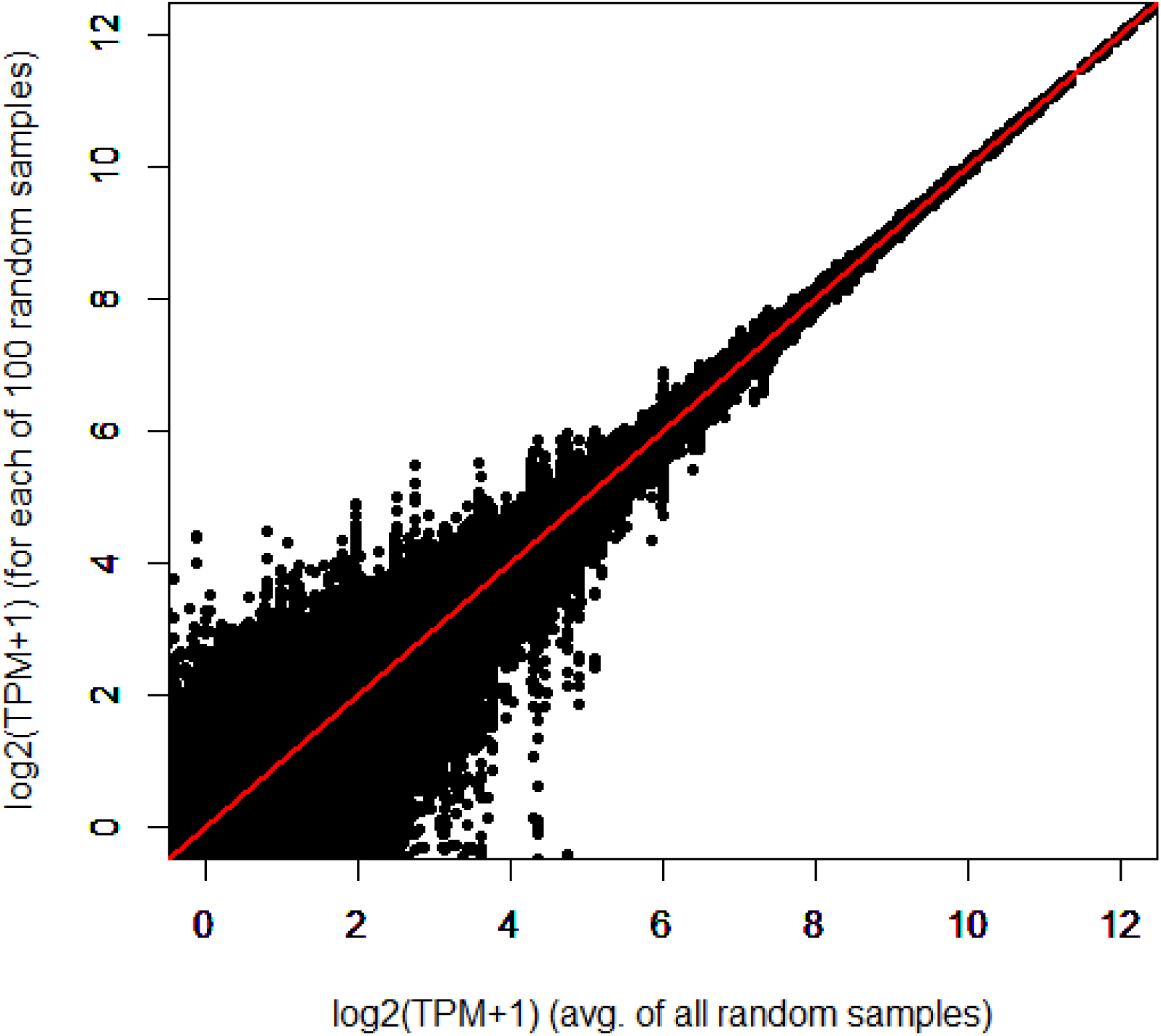
Randomly down-sampling RNA-seq reads has minimal impact on the overall expression profile, primarily affecting expression level estimates of lowly expressed genes. Data shown is from one dataset – unchallenged BMDMs from an adult female broiler (Ross 308) – although with quantitatively similar findings from other samples. The figure plots the average TPM per gene, taken after 100 random samples of 10 million reads, against the TPM obtained in each sample. The line *y* = *x* is shown in red.

### Biologically meaningful expression profiles are identified even after combining disparate RNA-seq datasets

If a meta-analytic approach to RNA-seq is valid, subsets of transcripts enriched in a given tissue should have annotations functionally appropriate to that tissue. To test this, we calculated a preferential expression measure (PEM) for each gene [65], essentially the median expression divided by the mean. We then obtained the set of Gene Ontology (GO) terms enriched in each subset of genes with the highest PEM associated with a particular tissue (Table S10) (see Materials and Methods). Consistent with the function of each tissue, the bursa of Fabricius (the site of B cell synthesis [66]) showed tissue-specificity for the expression of genes enriched for ‘defence response to bacterium’ (p = 8.3×10^−5^), breast muscle for ‘striated muscle contraction’ (p = 1.9×10^−6^), cerebrum for ‘synaptic transmission’ (p = 1.5×10^−4^), claw epithelium for ‘bone mineralisation’ (p = 6.4×10^−4^), heart for both ‘muscle contraction’ (p = 8.8×10^−6^) and ‘cellular respiration’ (p = 4.6×10^−15^), kidney for ‘oxidation-reduction process’ (p = 5.3×10^−5^), pancreas for ‘proteolysis’ (p = 0.001), pituitary gland for ‘endocrine system development’ (p = 2×10^−4^), retina for ‘visual perception’ (p = 7.2×10^−17^), spleen for ‘immune response’ (p = 2.2×10^−6^), and trachea for ‘cilium morphogenesis’ (p < 1×10^−30^) (Table S10). In an all-against-all correlation matrix (Pearson’s *r*) (Table S11), the expression profiles of like tissues were correlated regardless of their BioProject of origin (Table S12). A sample-to-sample network graph also demonstrates that samples of the same or related tissues cluster together (Figure 3). Taken together, these results validate the aggregation of data from multiple sources to create an informative expression atlas.

**Figure 3.**
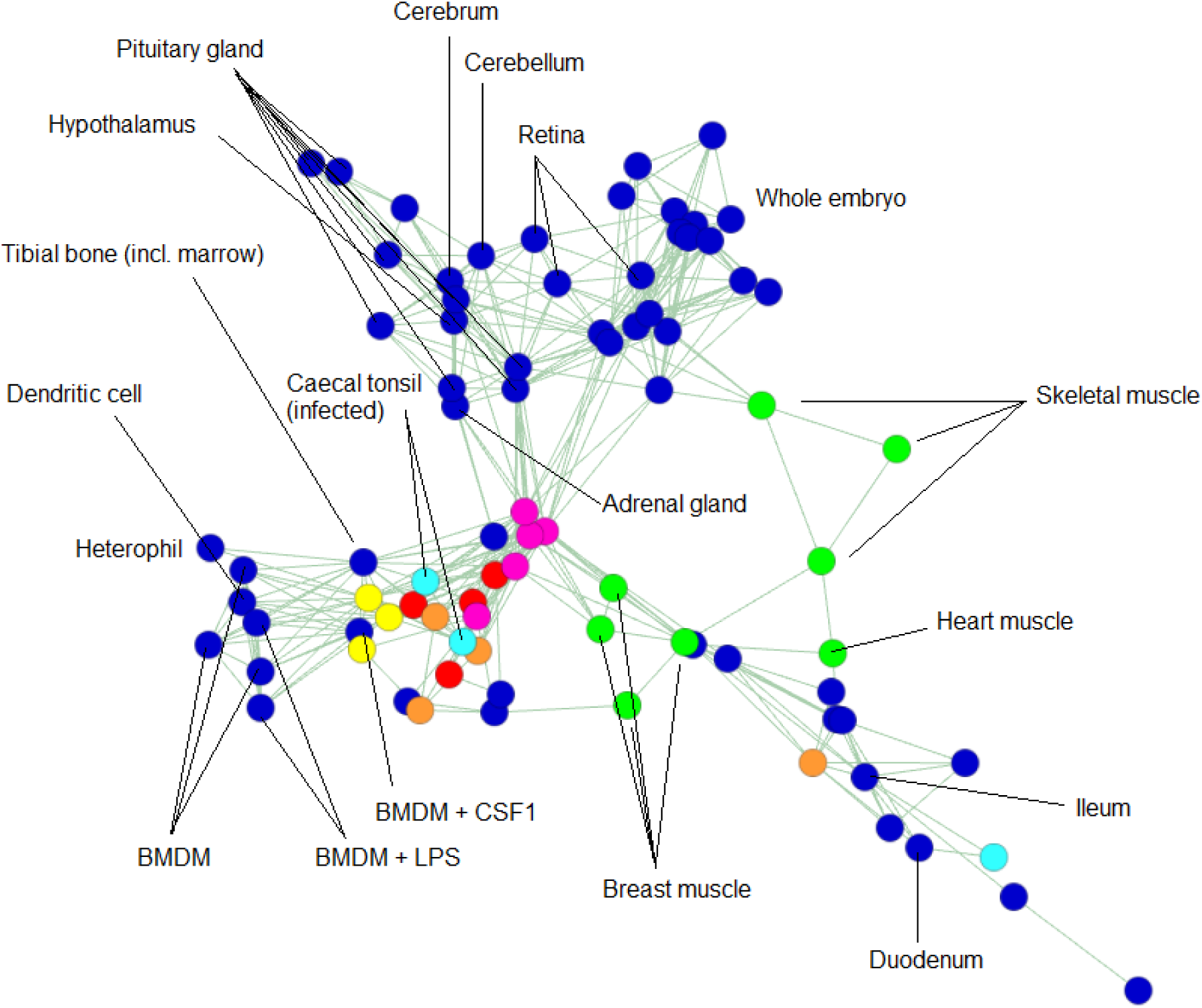
2D representation of a sample-to-sample network graph, plotting Spearman’s correlations between expression profiles. The graph was built using an RNA-seq meta-dataset with each sample distinct by tissue, developmental stage and BioProject of origin, and expression level per gene per sample averaged (where possible) across all replicates of that sample (dataset available as Table S6). Each node (circle) in the graph represents a sample, and each edge (line) a correlation exceeding a threshold (*rho* ≥ 0.82). The graph contains 82 nodes, connected by 243 edges. Selected nodes are labelled. Overall, like tissues tend to correlate more strongly with like, irrespective of BioProject of origin. Certain coloured nodes indicate tissues independently sequenced by multiple BioProjects (listed in Table S2), including liver (red), spleen (yellow), lung (orange), adipose (pink), caecal tonsil (light blue) and muscle (green). There are two notable idiosyncrasies: one of the four lung samples is comparatively dissimilar to the others of its group, as is one of the three caecal tonsil samples. In the latter case, however, the two most closely correlated caecal tonsil samples are those infected with *Campylobacter*. Consistent with this, these samples cluster more closely with immune cells and tissues. The third caecal tonsil sample belongs to a healthy chicken.

### Signals of co-expression allow for informative functional annotation

Network analysis of the meta-dataset was performed using Miru, a commercial version of BioLayout *Express*^3D^ [67, 68], previously applied to pig [26], sheep [27] and mouse [28] microarray datasets and CAGE data from the FANTOM5 consortium [24, 25]. A Pearson’s correlation matrix for each gene-to-gene comparison was visualised as a network graph of 18,127 nodes (genes) linked by 632,038 edges (correlations above a certain threshold; in this case, *r* = 0.8). Clusters of interconnected nodes represent sets of genes that share a signal of co-expression. These clusters were identified by applying the Markov clustering (MCL) algorithm [69] to the network graph, at an inflation value (which determines cluster granularity) of 2.2. The contents of each cluster are given in Table S13. Many of the co-expression clusters comprised genes with a tissue- or process-specific expression profile. Table S14 summarises the highest PEM value for a tissue in each of the clusters with >25 members. Cluster 2 was largely brain-specific: of the 655 genes in this cluster, 281 (43%) had their highest PEM in the hypothalamus, 155 (24%) had their highest PEM in the cerebrum and 115 (18%) had their highest PEM in the cerebellum. Other clusters contained genes with expression enriched in liver (cluster 6), ovary (cluster 7), trachea (cluster 8), testis (cluster 10), retina (clusters 13 and 24), feather epithelium (cluster 14), breast muscle (cluster 16), kidney (cluster 17), pituitary gland (clusters 19 and 25), *Campylobacter*-infected caecal tonsils (cluster 20), spleen (clusters 21 and 22) and adipose (cluster 23).

The tissues in some of these clusters were represented by multiple independent projects combined in this meta-atlas. For instance, cluster 6 comprises genes that were enriched in the liver, with data from three separate BioProjects. Some variation in expression estimates between these independent liver samples did not affect their inclusion in the same co-expression cluster. Furthermore, the GO terms enriched in each cluster are functionally consistent with its observed tissue-specificity (Table S15). Some clusters were associated with processes shared by multiple tissues. The largest cluster, cluster 1, was enriched in embryo-derived samples, and the GO terms are associated strongly with the cell division cycle and DNA repair (Table S15). The genes within this list include the key transcriptional regulator, *FOXM1*, and multiple cyclins (*CCNA2/B2B3/C/E1/F* and *J*), and overlap substantially with cell cycle-associated lists derived from previous cluster analysis [2, 70].

We used the ‘guilt by association’ principle to contextualise individual gene annotations – obtained by protein-level alignment and of varying quality (see Materials and Methods) – as there is an *a priori* expectation that by virtue of being co-expressed, the genes within a given cluster have related (that is, tissue- or process-specific) functions. In this respect, we can increase confidence in otherwise lower-quality alignments. Some examples and proposed annotations are summarised in Table S13. The co-expression profile is especially informative for clusters with few known genes. For instance, cluster 14 contains 210 genes expressed largely in the feather epithelium (Table S13). 93% of the genes within this cluster are unannotated, with only 14 genes having a known function (Table 1). Collectively, the functions of these genes are biologically consistent with an epithelium-enriched expression profile. Of the 196 unannotated genes, 86% can be aligned to feather keratins (representing 86 of the 96 genes with only an Ensembl ID and 83 of the 100 genes with only an NCBI RefSeq ID) (Table S13). Other unannotated genes include paralogues of existing genes in the cluster (ENSGALG00000004358 shares homology with *AMZ1*, ENSGALG00000029002 with *XG* and LOC428538 with *SDR16C5*), probable members of the keratin-associated protein family, which have essential roles in hair shaft formation [71] (ENSGALG00000018878, ENSGALG00000044257, LOC101751162, LOC101751279, LOC107055127, LOC107055128 and LOC107055130), a gene with homology to the tight junction protein claudin 4 (ENSGALG00000035131) [72], and several transcripts with homology to uricases (LOC101747367, LOC107056676 and LOC107056678), enzymes which degrade uric acid (the end point of purine metabolism) [73], notable because purines act as pigments in avian feathers [74].

**Table 1.** Genes in cluster 14 with known function.

| Gene symbol | Gene name | Protein function | References |
| --- | --- | --- | --- |
| AMZ1 | archaemetzincin 1 | metalloprotease, possibly involved in tissue remodelling to form feather follicles | [140, 141] |
| ANKK1<br>(PKK2) | ankyrin repeat and<br>kinase domain<br>containing 1 | interacts with keratin filaments | [142] |
| AREG | amphiregulin | epithelial growth factor | [143] |
| CLDN9 | claudin 9 | tight junction membrane protein found in all epithelia | [144] |
| DLX4 | homeobox protein<br>DLX4 | homeobox protein that regulates epithelial-mesenchymal interactions | [145] |
| EDMTF4<br>EDMTFH | epidermal<br>differentiation protein<br>starting with MTF<br>motif 4<br>epidermal<br>differentiation protein<br>starting with MTF and<br>rich in histidine | markers of the feather barbule and members of the epidermal differentiation complex;<br>this has a role in integumentary development, including feather pigmentation | [146-148] |
| FK21 | feather keratin 21 | feather keratins |  |
| FK27 | feather keratin 27 |  |  |
| PNPLA4 | patatin-like<br>phospholipase<br>domain-containing<br>protein 4 | enzyme with a role in retinol metabolism (retinol and related compounds regulate<br>epithelial cell growth and differentiation) | [149, 150] |
| RAB38 | Ras-related protein<br>RAB38 | GTPase involved in melanosome biogenesis and epithelial pigmentation | [151, 152] |
| RASSF10 | Ras association<br>domain family<br>member 10 | tumour suppressor that mediates the epithelial-mesenchymal transition | [153] |
| SDR16C5<br>(RDH-E2) | epidermal retinol<br>dehydrogenase 2 | overexpressed in psoriatic human skin | [154] |
| XG | Xg blood group | blood group antigen | [155] |

### Annotation of co-expression clusters associated with innate and acquired immunity and macrophage biology

The most prominent set of genes co-expressed in macrophages was cluster 4 (n = 458 genes; 129 [28%] are unannotated), in which > 60% of the genes have their highest PEM for BMDMs 24 hours post-LPS stimulation (Figure 4 and Table S14). This cluster is internally validated by the presence of transcripts encoding numerous known myeloid effectors/receptors (e.g. *C3AR1*, *CCR2*, *CD40*, *CYBB*, *CLEC5A*, *DCSTAMP*, *NLRC5*, *METRK*, *MYD88*, *TLR4*), lysosomal components (e.g. *CTSB*, *LAMP1*, *M6PR*) and multiple transcription factors (*BATF3*, *CEBPB*, *IRF1*, *NFE2L2*, *NRR1H3* [also known as *LXRA*], *SPI1* [also known as *PU.1*], *STAT1*, *TFEC*) that are also macrophage-enriched in mouse and human [75]. Their co-expression strongly indicates that basic macrophage transcriptional regulation is conserved between birds and mammals. Accordingly, the provisional annotations of genes that lack an informative name in this cluster, shown in Table S13, are given extra weight by their association. Other macrophage clusters include cluster 34 (n = 93 genes; 72 [77%] are unannotated) and cluster 37 (n = 79 genes; 16 [20%] are unannotated), in both of which the majority of genes had their highest PEM for the HD11 immortalised macrophage cell line (from BioProject PRJEB1406): 98% and 90%, respectively (Table S14). The smallest macrophage-specific cluster was cluster 84 (n = 26 genes; 19 [73%] are unannotated), in which every gene had its highest PEM for BMDMs treated with CSF1 (from BioProject PRJEB7662) (Table S14).

**Figure 4.**
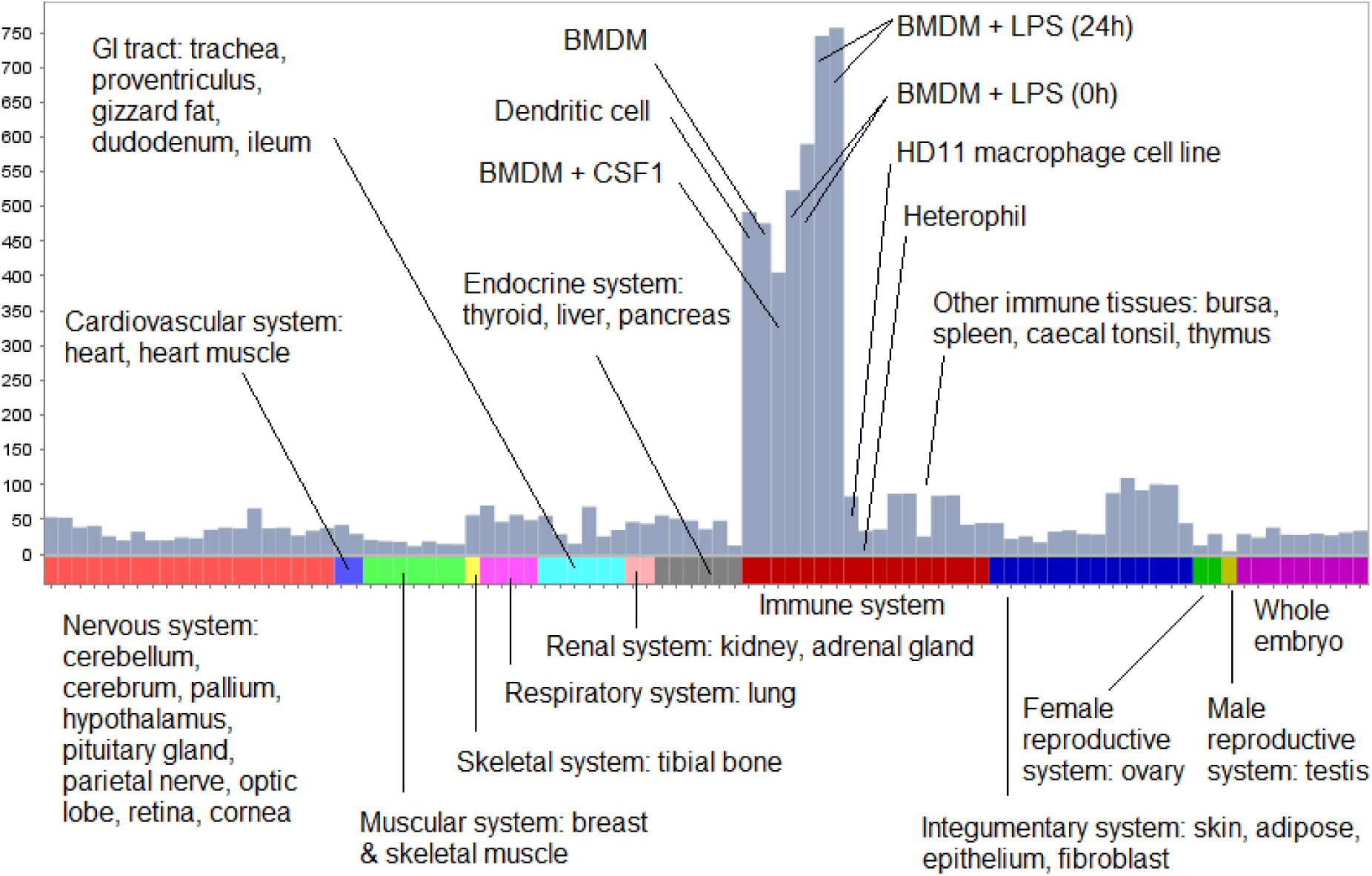
Expression profile of the macrophage-specific cluster 4. Histogram shows the average expression level of the 458 genes in the cluster, where expression level per gene is calculated as the median TPM across all replicates, per BioProject, per tissue. The expression level dataset is available as Table S6.

The *CSF1R* gene was contained within cluster 27 (n = 129 genes, of which 32 [25%] are unannotated), which had an expression profile shared by both dendritic cells and macrophages. 36% of the genes in cluster 27 had their highest PEM for dendritic cells and 26% for untreated BMDMs (both samples from BioProject PRJEB7475), with the remaining 26% for BMDMs treated with CSF1 (from BioProject PRJEB7662) (Table S14). This cluster also contained the lipopolysaccharide receptor and commonly used monocyte marker, *CD14*, several genes (*C1QA/B/C*, *MARCO*, *P2RY12/13*, and *STAB1*) that are associated with tissue-specific macrophage populations in mice [76], and a single myeloid-associated transcription factor, *MAFB*, which is required for tissue macrophage development in mice [77]. The cells referred to as dendritic cells are bone marrow cells grown in *GM-CSF* (*CSF2*), rather than *CSF1*. As noted in previous analyses of mouse [78] and human [79] transcriptomes, cells differentiated in *GM-CSF* have much more in common with macrophages than with classical dendritic cells dependent upon *FLT3*-ligand.

The clusters associated with the acquired immune response, predominantly B and T cells, are somewhat smaller and poorly-annotated (clusters 20, 21, 22, 29 and 78). Cluster 21, expressed most highly in spleen, contains *TIMD4* (ENSGALG00000003876), which promotes T-cell expansion and survival [80], and is enriched with B cell-associated genes, including the B cell transcription factors *BATF*, *IRF4*, *PAX5*, *RUNX3*, and *SPIC*, as well as the class II trans activator *CIITA*, class II subunit *CD74* and the class II MHC gene *BLB2*. The thymus-enriched cluster 29 contains *CD4*, the recombination activating genes *RAG1* and *RAG2*, and the T cell transcription factors *LEF1*, *RORC* and *TCF7*.

### Integrating gene expression and protein-protein interaction networks

Biological systems can be functionally organised into many different (and intersecting) networks based on the nature of their interaction, including – aside from gene co-expression networks – metabolic/biochemical networks, signal transduction networks, regulatory networks, and protein-protein interaction (PPI) networks [81]. Data from different networks can be integrated: for instance, subunits of the same protein complex are known to be co-expressed [82], with those genes present in both a co-expression and PPI network having a high probability of performing similar functions [83]. We therefore determined the set of genes present in both the same co-expression cluster and a PPI network (Table S16), obtaining chicken PPI data by mapping human PPIs to orthologous chicken genes (see Materials and Methods). The PPI and co-expression data are mutually supportive. For example, there were 32 PPIs among the genes in the macrophage-specific cluster 4. These include *STAT1* (signal transducer and activator of transcription 1-alpha/beta) – a critical mediator of the pro-inflammatory response of macrophages to LPS [84] – and the transcription factors *ATF3*, a known inducer of *STAT1* [85], and *SPI1/PU.1*, which is essential for macrophage differentiation [86]. Also in the network are the tyrosine kinase *LYN*, which is activated alongside *STAT1* in response to *IL5* (a key mediator of eosinophil activation [87]), and the adaptor protein *GRB2*, which facilitates the activation of *ERK* by tyrosine kinases [88] (*ERK* signalling is essential to macrophage development [89]). In addition, the network contained *SOCS3*, a negative regulator of cytokine signalling that inhibits the nuclear translocation of *STAT1* in response to *IFN* stimulation [90], with this stimulation a key constituent of classical macrophage activation [91].

### Integrating gene expression and promoter expression networks

Relatively few RNA-seq datasets were available for chicken embryonic development. Lizio *et al* [53] have recently analysed the time course of chicken development using Cap Analysis of Gene Expression (CAGE). Their dataset complements a CAGE-based analysis of gene expression in multiple tissues of the mouse during embryonic development [92]. Network analysis of the mouse dataset revealed a signature of the expansion of the tissue macrophage populations during embryonic development, and the inverse relationship between cell proliferation and tissue-specific differentiation in each organ [93]. Analysis of a macrophage-specific transgene in birds revealed that, as in mammals, macrophages are first produced by the yolk sac, progressively infiltrate the embryo and expand in number to become a major cell population in every organ [33, 94]. The expression atlas we have developed provides a complementary resource for adult tissues and includes a time course of embryonic development. By combining the atlas with the CAGE data, it would be possible to infer the developmental time course of organ systems in the chicken. We obtained the chicken CAGE data of Lizio *et al* [53] and clustered the promoter-based expression levels in the same manner as for the RNA-seq atlas. Figure 5 shows the resulting network graph, and the average expression profiles of a subset of clusters. Table S17 provides a full list of promoters in each of the co-expression clusters and their average expression profiles. As discussed by Lizio, *et al*, the embryonic CAGE data identify transcription start sites for many tissue-specific and regulated genes, including developmental regulators such as *brachyury*. The intersection of the CAGE and RNA-seq clusters is presented in Table S18. Not surprisingly, the largest promoter cluster overlapped substantially with cluster 1 in the RNA-seq atlas which was embryo-enriched in expression. It contained numerous developmental regulators, anabolic/cell cycle, and mitochondria-associated genes with an average profile of down regulation during development (Figure 5). Aside from the whole embryo profiles, the CAGE data contains several additional samples, including bone marrow-derived mesenchymal stem cells (MSC), aortic smooth muscle cells (ASMC), hepatocytes, extra-embryonic tissues and both leg and wing buds. Each of the samples was enriched for specific promoters that also varied during development and accordingly defined clusters. Clusters 2 and 10 of the CAGE data were enriched in MSC and ASMC, and contained many mesenchyme-associated genes including multiple collagens and other connective tissue-associated transcripts. CAGE clusters 4 and 9 were hepatocyte-enriched and most likely track the development of the liver during development. Cluster 4, shown in Figure 5, contains the transcription factor *HNF1A*, and many of the transcripts within it encode secreted proteins such as complement components and clotting factors. CAGE cluster 5 (Figure 5) contains the muscle-specific transcription factors *MYOD1*, *MYOG* and *SOX2*, and numerous skeletal muscle-associated genes in common with cluster 16 from the RNA-seq atlas, and increases in expression throughout development. The transcripts within cluster 5 are not expressed in the aortic smooth muscle cells. CAGE clusters 7, 16, 18 and 19 contained transcripts that were expressed transiently at different stages of embryonic development, including multiple members of the *HOX* and *CDX* families. CAGE clusters 8 and 25 both contained promoters of multiple genes that are expressed specifically in macrophages in the RNA-seq atlas (clusters 4 and 27). The average expression profiles are shown in Figure 5, with representative genes indicated. The macrophage-specific transcription factor *SPI1*, and most other macrophage-enriched genes within CAGE clusters 8 and 25, fall within the larger macrophage-associated clusters (4, 27 and 31) within the RNA-seq atlas. Interestingly, CAGE cluster 25 appears to be enriched for genes expressed specifically in brain macrophages (microglia), including *CSF1R*, *C1QA, C1QB, C1QC, CTSS, DOCK2, HAVCR1 LAPTM5, LY86, MPEG1*, and *P2RY13* [95], which in mice appear to develop from yolk sac progenitors rather than definitive haematopoiesis [96]. Several other microglia/macrophage associated transcripts, notably *CX3CR1, P2RY12, TIMD4*, and *TREM2*, are detectable in the CAGE data at the same embryonic stage, but did not cluster because their expression differs in the cell populations. In each of the macrophage-associated clusters, there were numerous promoters currently with uninformative annotation, which by inference are likely to be macrophage-related. Consistent with the location of *CSF1R* mRNA and the *CSF1R*-reporter gene in the chicken [33], *CSF1R* and *SPI1* were both first detectable in the embryo at between HH12 and HH14 (day 2), and both increased in parallel during embryonic development. Figure 6 shows the ZENBU (http://fantom.gsc.riken.jp/zenbu/) view of the chicken *CSF1R* locus, identifying the transcription start site downstream of the *PDGFRB* locus, and the time course of appearance of *CSF1R* transcripts in the embryo and their expression in isolated cells. The reason that CAGE clusters 8 and 25 genes separate in the dataset is that they were also detected at high levels in “mesenchymal stem cells” and to a varying extent in “hepatocytes” (Figure 5). In mice, macrophages were shown to be a major contaminant of bone marrow-derived osteoblast cell cultures [97]. Based upon this cluster analysis in the embryo (which reveals separate mesenchyme and hepatocyte-specific clusters), and the atlas data, where these genes were clearly macrophage-enriched, the expression of macrophage-associated genes is almost certainly a reflection of the presence of large numbers of macrophages in these cell populations. Indeed, the set of promoters active in “mesenchymal stem cells” was found to be enriched for binding sites for *SPI1* and *CEBPA*, transcription factors that can induce the transdifferentiation of lymphoid precursors into macrophages [98].

**Figure 5.**
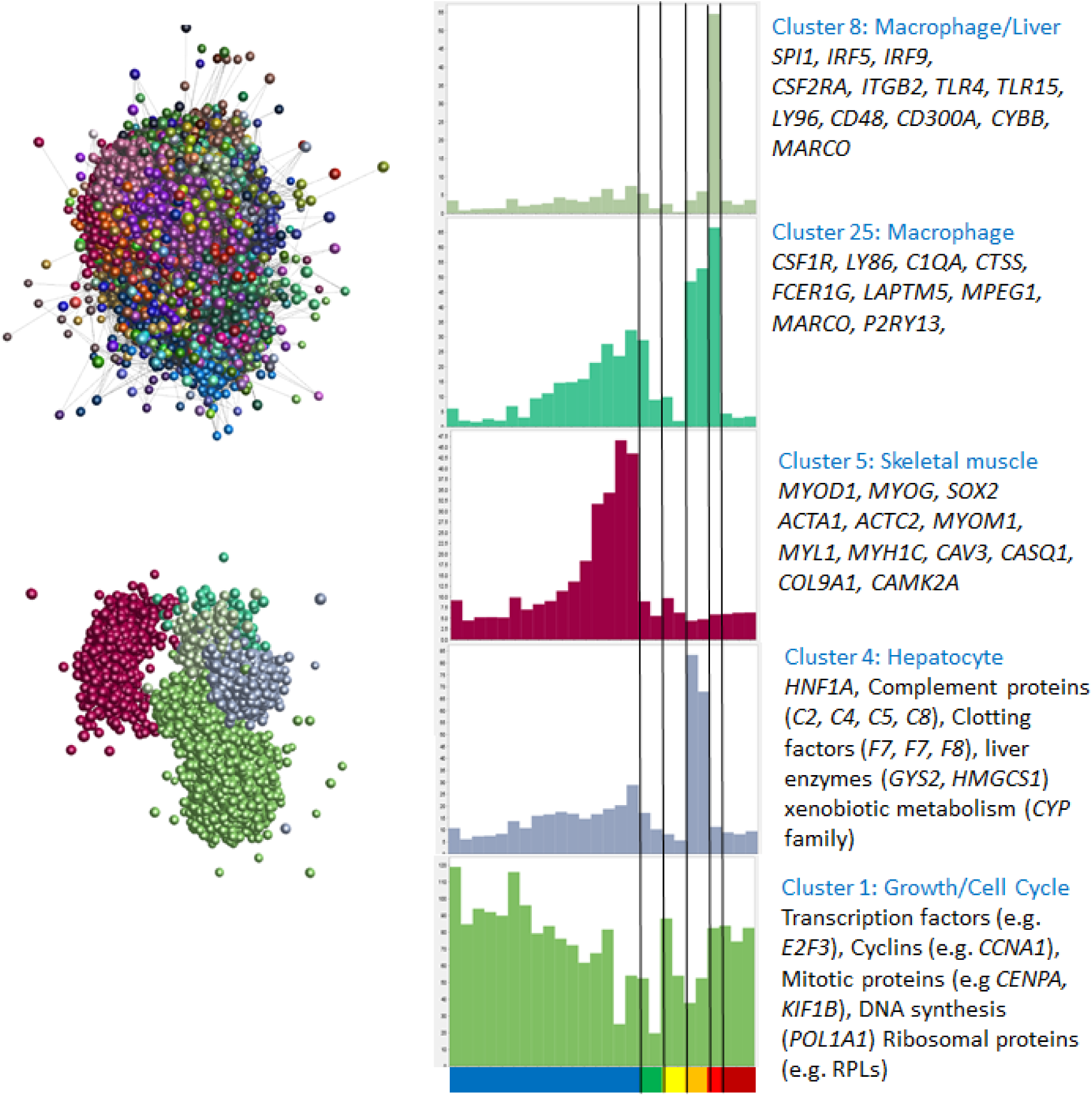
The panel on the left shows the clustered nodes for the main element in the layout graph (upper section). Nodes allocated to the same cluster are the same colour. The panel on the right shows the average expression profiles for five clusters highlighting the different phases of chick embryo development, and key genes for each cluster are shown in the boxes. The layout of these clusters within the main element is shown in the lower part of the left panel. Node colour matches the colour of the bars on the histograms. The X-axis shows the different samples (blue – embryo developmental time course from 1.5 hours to day 20 after fertilisation (HH45); green – extraembryonic tissues; yellow – limb buds; orange – hepatocytes; red – bone marrow derived mesenchymal stem cells; dark red – aortic smooth muscle cells. Full detail of the samples can be found in Lizio, *et al* [53]. Y axis shows average TPM for TSS in the cluster for each sample.

**Figure 6.**
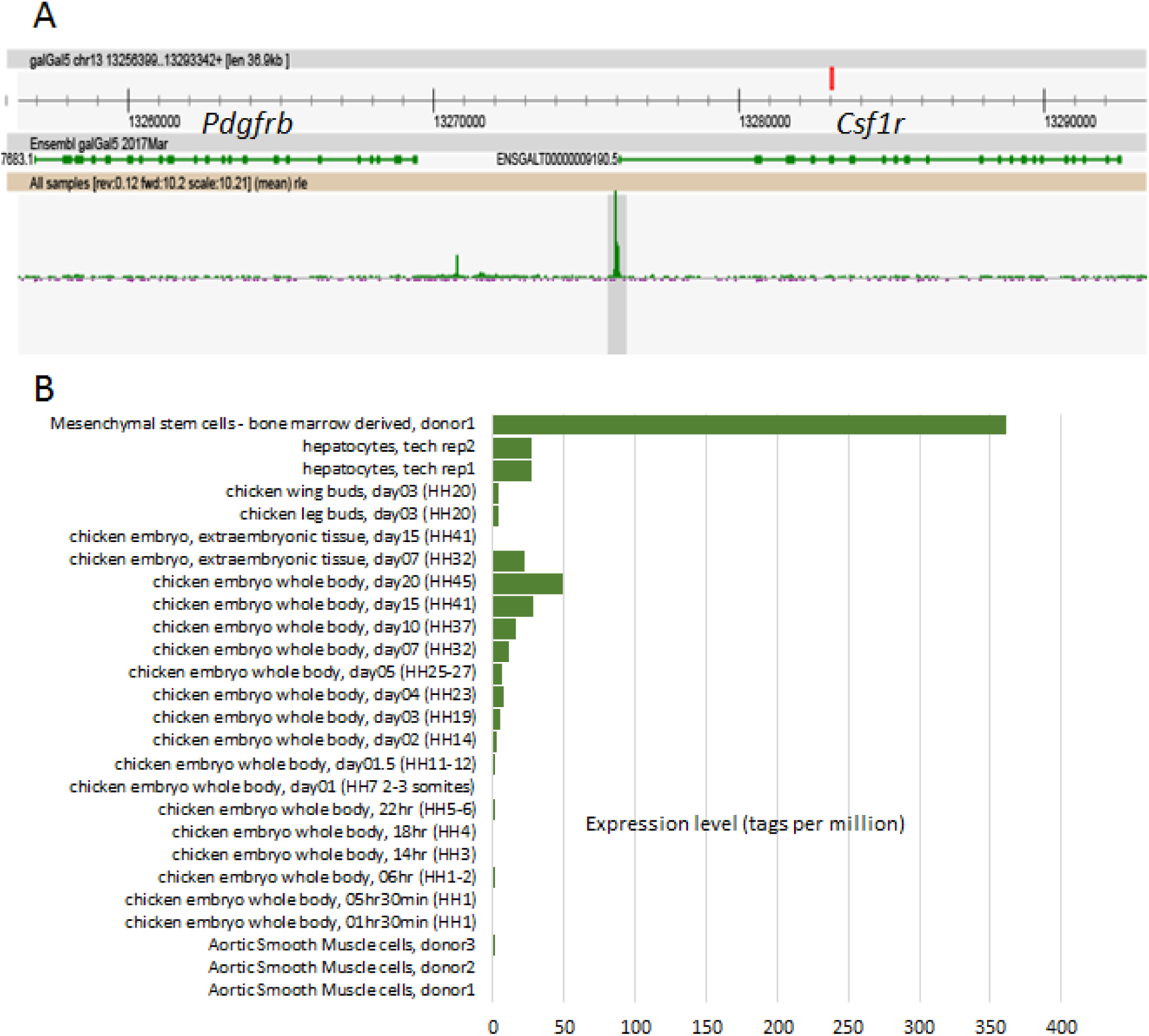
ZENBU (http://fantom.gsc.riken.jp/zenbu/) view of the chicken *CSF1R* locus, identifying the transcription start site downstream of the *PDGFRB* locus (A), and the time course of appearance of *CSF1R* transcripts in the embryo and their expression in isolated cells (B).

## DISCUSSION

RNA-seq is a multi-step process of reverse transcription, amplification, fragmentation, purification, adaptor ligation and sequencing, with each step subject to error [99]. Such laboratory-specific variation is also independent of intrinsic sequencing biases, which can influence the nucleotide composition of the reads [100] (leading to mismatches between the sequenced read and the original RNA fragment [101]), the GC content of the reads [102], and the sequencing error rate [11]. Despite all of these constraints, Figure 3 shows that in a sample-to-sample network graph of many independently sequenced tissues, the signal of co-expression clearly outweighs the noise.

The critical step in reducing the noise, and making the datasets comparable, was to down-size the RNA-seq libraries so that the depth of coverage of the transcriptome was the same in each case. This has the effect of removing a great deal of the stochastic detection of more lowly-expressed transcripts. Figure 2 and Table S9 show that the random sampling used to down-size does not substantially alter the relative expression estimates of any two genes within any given sample, with equivalent expression profiles reconstructed for each of 100 random samples. Combined with the use of Kallisto to quantify expression, which maps a common depth of k-mers to a standardised reference transcriptome, the method we have developed effectively ensured that each RNA-seq library was exploring an equivalent transcriptomic space.

The success of the aggregation of public domain data in terms of genome annotation is evident from the analysis of the membership of co-expression clusters in Table S13. Each cluster clearly contains genes of known function, shows evidence of very strong GO enrichment, and as noted in similar array-based studies [2, 26] commonly contains the transcription factors that regulate the other members of the cluster. On that basis, it would be reasonable to provisionally assign the same GO terms to genes of unknown function, at least within the larger clusters. For example, the genes within cluster 1 that are not currently functionally annotated or assigned a clear orthologue are likely to be involved in some way in the cell cycle. Indeed, the provisional annotations of many of them shown in Table S13 indicate this is very likely to be the case. Similarly, the genes we have identified that were enriched in innate and acquired immune cells are likely to be associated with heritable variation in disease resistance/susceptibility.

Detailed examination of individual clusters can provide significant biological insights. Cluster 8, enriched in trachea, and with the second highest expression in lung, was strongly enriched with GO terms associated with cilium, microtubule binding, motor activity and the actin cytoskeleton (Table S15), and includes, for example, multiple members of the cilia and flagella-associated protein (*CFAP*), dynein regulatory complex (*DRC*) and other dynein-related gene families. Mutations in many of these genes have been associated with human ciliopathies [103]. This cluster also contained the transcription factor *FOXJ1*, which is essential for the formation of motile cilia in mice [104]. Provisional annotations of genes of unknown function in this cluster are consistent with the overall enrichment for genes associated with motility. The presence of the epithelial transcription factors *ELF5* and *PAX9* in this cluster suggests both could have a role in regulation of this key gene set, providing a possible reason for the embryonic lethality of the knockouts of each gene [105, 106]. Interestingly, *KIAA0586*, which is also known as *TALPID3*, is in a separate smaller cluster – number 139 – that is more widely expressed. The *TALPID3* protein encodes a centromeric component, and mutation affects the formation of primary, non-motile cilia and signaling by the morphogen sonic hedgehog [107, 108]. Many of the genes that are apparently co-regulated with *TALPID3* have been associated in some way with regulatory functions of primary cilia, including *CEP120* which, like *KIAA0596*, is mutated in human Joubert syndrome [109]. Other members of the cluster may be candidate interactors with *TALPID3*.

The validity of the approach, and of the clusters generated, was established by comparing tissue- and function-specific clusters obtained by an alternate method of quantifying RNA expression levels, CAGE, using a public dataset of chicken embryo development. This showed that tissue-specific developmental gene expression can be detected using whole embryos (as we have previously shown for mouse [93]), and that the genes in the developmental stage clusters matched those found in the adult tissue atlas.

The clustering we have presented is based upon an arbitrary correlation threshold. For every gene of interest, it can be informative to identify its transcriptional companions. To this end, as we have done previously for human [2], pig [26], sheep [27] and mouse [28], we have made the current version of this atlas available as a searchable database using the gene annotation portal BioGPS [110] (http://www.biogps.org/chickenatlas), where one can utilise a simple “find correlated” function to identify genes with similar expression profiles. In turn, this resource allows a rapid comparative assessment of the expression of a gene of interest in mammals and birds and the extent to which functional information is likely to be transferable across species.

The advantage of the aggregation method we have applied is that it is can be extended with new data from tissues and cell types we have not currently included. The larger the dataset, and the greater the transcriptional space sampled, the more stringent the correlations that will be generated and the more likely they are to produce new biological insights.

## MATERIALS AND METHODS

### Animals

To obtain bone marrow-derived macrophages, nine chickens of approximately 8 weeks of age (3 female and 3 male Ross 308 broilers, and 3 female CSF1R-MacApple transgenic NOVOgen Brown layers) were euthanized by cervical dislocation and confirmed dead by decapitation. Likewise were euthanized 23 broiler chickens, each 5 weeks of age, to obtain the caecal tonsils. All animal work was conducted in accordance with guidelines of the Roslin Institute and the University of Edinburgh and carried out under the regulations of the Animals (Scientific Procedures) Act 1986. Approval was obtained from the Roslin Institute’s and the University of Edinburgh’s Protocols and Ethics Committees.

### Macrophage cell culture and RNA isolation

Bone marrow-derived macrophage (BMDM) culture and challenge *in vivo* were performed as previously described [111]. Chicken bone marrow was cultured for 7 days with 350 ng/µl chicken CSF1 on Sterilin plastic to differentiate BMDMs. Adherent cells were then transferred to tissue culture plastic and cells plated at 80% confluence. BMDMs were challenged with the addition of LPS at 100 ng/ml to culture medium and then harvested after 0 (null condition), and 24 hours. Cells were harvested in TRIzol^®^ (15596018; Thermo Fisher Scientific) and extraction performed with the RNeasy Mini Kit (74106; Qiagen Hilden, Germany) according to manufacturer’s instructions.

### Collection of Campylobacter-infected caecal tonsils

Birds were naturally exposed to *Campylobacter* spp. under commercial farm conditions. Caeca and caecal tonsil samples were collected in RNAlater (AM7021; Thermo Fisher Scientific, Waltham, USA). *Campylobacter* load in caeca was determined by selective culture as previously described [112]. Seven serial ten-fold dilutions of caecal content were prepared in phosphate-buffered saline and 100 μl plated to mCCDA (modified cefoperazone-deoxycholate agar) supplemented with cefoperazone (32 mg/L) and amphotericin B (10 mg/L; Oxoid), followed by incubation for 48 hours under microaerophilic conditions (5% O2, 5% CO2, and 90% N2) at 41C. Dilutions were plated in duplicate and colonies with morphology typical of *Campylobacter* detected in all samples. RNA was extracted from the caecal tonsils using the RNeasy Mini Kit (74106; Qiagen Hilden, Germany) according to manufacturer’s instructions. As chickens were exposed naturally rather than being explicitly challenged with *Campylobacter*, bacterial load varied considerably between individuals. Accordingly, tonsil samples were partitioned into two broad subsets: those from chickens whose caecum has high *Campylobacter* load (>= 10,000 CFU/g), and those with low *Campylobacter* load (< 10,000 CFU/g).

### RNA-sequencing

For both BMDM and caecal tonsil samples, library preparation was performed by Edinburgh Genomics. Total RNA (for BMDMs) and mRNA (for caecal tonsils) was, in both cases, sequenced by Edinburgh Genomics at a depth of >40 million strand-specific 75bp paired-end reads per sample, using an Illumina HiSeq 4000. The raw data is deposited in the European Nucleotide Archive under accessions PRJEB22373 (BMDMs) and PRJEB22580 (caecal tonsils).

### Public RNA-seq datasets

Publicly accessible datasets used in this study are described in Table S1. The meta-atlas aggregating these data details, per tissue, the associated NCBI BioProject and Sequence Read Archive (SRA) sample IDs (Table S6). All public datasets for this study are available via the SRA, a public repository for sequence data maintained by the International Nucleotide Sequence Database Collaboration (INSDC) and accessible from the websites of its constituent members: known as the SRA if via the National Center for Biotechnology Information (NCBI) (www.ncbi.nlm.nih.gov/sra), the DRA (DDBJ Read Archive) if via the DNA Data Bank of Japan (DDBJ) (http://trace.ddbj.nig.ac.jp/dra/), and the European Nucleotide Archive (ENA) if via the European Bioinformatics Institute (EBI) (www.ebi.ac.uk/ena) [113]. For retrieving the raw files used in this study or for expanding this work with new datasets from novel tissues, note that data are directly accessible in fastq format from the ENA and DDBJ but only in a binary.sra format from the NCBI. Decompiling the latter into fastq files – using the fastq-dump tool within the SRA Toolkit (https://trace.ncbi.nlm.nih.gov/Traces/sra/sra.cgi?cmd=show&f=software&m=software&s=software) – is far slower than analysing fastq files with Kallisto, and so forms a bottleneck in the expression atlas creation pipeline. For this reason, obtaining fastq files in bulk from NCBI is not recommended unless necessary.

### Defining a reference transcriptome and quantifying expression

Prior to expression level quantification, all RNA-seq datasets were randomly down-sampled to 10 million reads using seqtk (https://github.com/lh3/seqtk, downloaded 29th November 2016) with parameter -s 100 (to seed the random number generator). Expression level was then estimated, as transcripts per million (TPM), using the high-speed quantification tool Kallisto v0.43.1 [60] and default parameters. For datasets comprising single-end reads, we used parameters -l 100 -s 10; estimates of the average fragment length and standard deviation of the fragment length, respectively. Kallisto quantifies expression at the transcript level by building an index of k-mers from a set of reference transcripts and then mapping the RNA-seq reads to it, matching k-mers generated from the reads with the k-mers present in the index. Transcript-level TPM estimates are then summarised to the gene level. A critical aspect of this method is in selecting an appropriate set of reference transcripts for which expression is quantified. An appropriate value of *k* for the index is also required because if *k* is too large relative to read length, there is a higher chance the k-mers of the reads will contain errors (as read quality decreases towards the 3’ end of reads [4]). If the reads generate erroneous k-mers, they will not match the k-mers of the index. We used a value of *k* = 21, which lies – approximately – between half the length of the shortest read and a third the length of the longest read.

As a reference transcriptome, we obtained from Ensembl v89 the set of GalGal5 protein-coding transcripts, parsing the batch release (ftp://ftp.ensembl.org/pub/release-89/fasta/gallus_gallus/cds/Gallus_gallus.Gallus_gallus-5.0.cds.all.fa.gz, accessed 21^st^ June 2017) to retain only those transcripts with the ‘protein-coding’ biotype (n=28,768 transcripts, representing 10,846 genes). To this was added the CDS of 28,466 NCBI mRNA RefSeqs that had neither been assigned Ensembl transcript IDs, nor whose sequence was already present in the Ensembl release (under any other identifier). To reduce the likelihood of spurious read mapping, CDS < 300 bp were excluded from analysis. Erroneous expression level estimates are more likely when fewer possible reads can be derived from a gene, i.e. if the CDS is short [59]. While this approach arguably improves accuracy, it unavoidably excludes certain families, for instance the gallinacins [114], antimicrobial peptides known for their short chain lengths [115].

Although the Ensembl and NCBI sets of transcripts overlap, there are many unique entries in each. For example, RefSeqs XM_015294055 and XM_015294059 are both predicted transcripts of the macrophage-marker gene *CD163* [116], although Ensembl refers to this gene only by the numerical ID ‘418303’. RefSeq records beginning ‘XM’ are produced by the NCBI genome annotation pipeline and can lack transcript or protein homology support; by contrast, ‘NM’ records are validated [117]. Consequently, neither of the *CD163* RefSeqs are assigned Ensembl transcript IDs, and so they are excluded from the Ensembl batch release.

The RefSeq mRNA set also includes predictions of novel transcript sequences for existing Ensembl genes. For instance, the chicken *BF1* gene (classical MHC class 1; Ensembl gene ID ENSGALG00000033932) has 7 transcripts (Ensembl v89), encoding proteins of length 228, 323, 345, 346, 350, 354 and 360 amino acids (aa). However, *BF1* has only 3 associated mRNA RefSeqs, 1 validated and 2 predicted: NM_001044683, XM_015294995, and XM_015294996. These RefSeqs do not necessarily encode different proteins to those present in Ensembl – rather, the RefSeq mRNAs incorporate untranslated regions (UTRs) and so can encapsulate Ensembl CDS. For instance, the validated RefSeq mRNA NM_001044683 encodes the same 360aa protein as Ensembl CDS ENSGALT00000066783 (i.e. the same transcript model is independently available from both resources), but the RefSeq nucleotide sequence extends 17 bases upstream (the 5’ UTR) and 146 bases downstream (the 3’ UTR) of the coding ORF. By contrast, XM_015294995 encodes a putative 356aa peptide (XP_015150481) and XM_015294996 a 349aa peptide (XP_015150482), neither of which are available from Ensembl. As the XM_015294996 mRNA – an automated prediction – fully incorporates ENSGALT00000086848 (the CDS encoding the 228aa *BF1* protein), we considered the sequence better supported by the Ensembl model, as Ensembl takes a conservative approach to annotation [118], and the predicted peptide spurious. By contrast, the XM_015294995 mRNA does not contain any existing Ensembl CDS and so encodes a protein absent from Ensembl.

Overall, we retained RefSeq ‘XM’ mRNAs only if they can be assigned to a gene not yet present in the Ensembl annotation, or, if that gene is present, they do not incorporate a CDS from any of that gene’s Ensembl transcript models. UTRs were trimmed from each RefSeq mRNA by excluding all sequence outside the longest ORF. This combined set of Ensembl and RefSeq transcripts constitutes a standardised RNA space against which expression can be quantified, as in [59].

After quantifying expression with this initial transcriptome, a revised transcriptome was created, excluding those transcripts whose average TPM was < 1 in all tissues (Table S5), or which were only detectable in one tissue (as these may be artefacts of differential sequencing depth). Tissues whose distribution of TPM estimates does not comply with Zipf’s law (see below) were not counted. The revised transcriptome contains 28,276 Ensembl transcripts (representing 10,826 Ensembl genes) and 26,694 NCBI transcripts (which account for only 4665 existing Ensembl genes).

### Compliance of RNA-seq datasets with Zipf’s law

In a correctly prepared RNA-seq dataset, a minority of reads will produce the majority of reads and so its distribution of gene-level TPM estimates should comply, to a reasonable approximation, with Zipf’s law (which states that the probability of an observation is inversely proportional to its rank). A custom Perl script was used to identify, per sample, the number of unique TPM values and the number of genes with a TPM at or exceeding this level. After excluding, for robustness, data from the first and last order of magnitude (as in [119]) and all values of TPM < 5 (which have a higher likelihood of transcriptional noise), the data was log-transformed and a linear regression model fitted using R v3.2.0 [120]. Samples whose exponents deviated too greatly from -1 (by ± 20%, i.e. if the exponent is < -0.8 or > -1.2) were considered erroneous.

### Tissue specificity

For each gene, we calculated a preferential expression measure (PEM) in a manner similar to [65]. PEM relates the average expression of that gene in a given tissue to the average expression of that gene in all tissues. For each gene *i*, then for tissue *t*^*i*^, PEM(*t*^*i*^) = S-A, where S = expression of gene *i* in tissue *t*^*i*^, and A = arithmetic mean expression of gene *i* across the set of all tissues. Prior to calculation, all TPM values < 1 were considered to be 1, and a log^2^-transformation applied. This is to ensure that genes with expression indistinguishable from noise (TPM < 1) will have a PEM of 0. Each gene will have a distribution of PEM values, one for each tissue in the meta-datasets. Genes with higher PEM values for a given tissue are more tissue-specific in their expression profile.

### Gene Ontology (GO) term enrichment

GO term enrichment was assessed using the R package topGO [121], which utilises the ‘weight’ algorithm to account for the nested structure of the GO tree [122]. topGO requires a reference set of GO terms, which was built manually from the GalGal5 set (obtained from Ensembl BioMart v89 [123]) and filtered to remove those terms with evidence codes NAS (non-traceable author statement) or ND (no biological data available), and those assigned to fewer than 10 genes in total Significantly enriched GO terms (p < 0.05) are reported only if the observed number per tissue exceeds the expected by 2-fold or greater.

### Gene annotation

Unannotated genes in GalGal5 – those with only an Ensembl placeholder ID, rather than an HGNC name – are annotated by reference to the NCBI non-redundant (nr) peptide database v77 [124], with each annotation assigned a quality category of 1 to 8 (highest to lowest quality, respectively), as previously described [27]. For each unannotated gene, we took the longest encoded peptide and obtained the set of blastp alignments [125] against NCBI nr, at a scoring threshold of p <= 1e^−25^. These alignments are a set of possible gene descriptions, of which only one can be selected as the annotation of that gene. The lowest quality category, 8, is the blastp hit with the lowest E-value. All subsequent quality categories require higher-quality hits, which: (a) have a % identity within the aligned region of >= 90%, (b) have an alignment length >= 90% of the length of the query protein, (c) have an alignment length >= 50 amino acids, (d) have no gaps, and (e) are not to a protein labelled either ‘low quality’, ‘hypothetical’, ‘unnamed’, ‘uncharacterized’ or ‘putative’, or otherwise having a third-party annotation (as these can be by inference and not experiment). Quality category 7 is the best-scoring (i.e. lowest E-value) of these higher quality hits. Category 6 is as above, but with at least one identifiable hit to the human proteome. Category 5 requires that the set of alignments span at least 4 different genera (excluding *Gallus*). At this point, if >= 75% of the alignments have the same description, the gene is named for the associated HGNC name (according to ftp://ftp.ebi.ac.uk/pub/databases/genenames/new/tsv/locus_types/gene_with_protein_product.txt, downloaded 24^th^ August 2016). However, as NCBI nr aggregates multiple sources of data, gene descriptions have numerous synonyms and so it is not always possible to automatically assign an HGNC symbol. The highest quality categories, 1 to 4, not only meet the above criteria but have degrees of reciprocal % identity to the human proteome. The highest quality category, 1, is if there is also a near-perfect match to an existing, related, peptide (alignment length >= 90% of the length of a human protein). Other quality categories, in descending order, are: 2 (alignment length >= 75% of the length of a human protein), 3 (>= 50%), and 4 (< 50%). Human protein sequences were obtained from genebuild GRCh38.p8 (ftp://ftp.ncbi.nlm.nih.gov/genomes/all/GCF_000001405.34_GRCh38.p8/GCF_000001405.34_GRCh38.p8_protein.faa.gz, downloaded 30^th^ August 2016).

### Network analysis

Network analysis was performed using Miru (Kajeka Ltd, Edinburgh, UK), a commercial version of BioLayout *Express*^3D^ [67, 68]. Miru determines the similarities between individual expression profiles by building a correlation matrix for both gene-to-gene and sample-to-sample comparisons. This matrix is then filtered to remove all correlations below a certain threshold (for the gene-to-gene comparison in the RNA-seq atlas, Pearson’s *r* < 0.8). A network graph is constructed by connecting nodes (genes) with edges (correlations above the threshold), and its local structure interpreted by applying the Markov clustering (MCL) algorithm [69] at an inflation value (which determines cluster granularity) of 2.2.

### Protein-protein interactions

Protein-protein interaction data was obtained from the IID (Integrated Interactions Database) version 2017-04 (http://iid.ophid.utoronto.ca/iid, accessed 25th July 2017) [126], a resource which combines computationally predicted PPIs with experimentally determined PPIs drawn from multiple databases. These include BIND (Biomolecular Interaction Network Database) [127], BioGRID (Biological General Repository for Interaction Datasets) [128], DIP (Database of Interacting Proteins) [129], HPRD (Human Protein Reference Database) [130], IntAct [131], I2D (Interologous Interaction Database) [132], InnateDB [133] and MINT (Molecular Interaction Database) [134]. The format of the PPI data is as a list of UniProt IDs, with one of three evidence types for the interaction: ‘exp’ (experimentally determined in this species), ‘pred’ (an *in silico* prediction from one of four previous studies [135–138]) and ‘ortho’ (predicted by mapping experimentally determined PPIs from another species to orthologous protein pairs in this species). As chicken PPI data is unavailable, we obtained human PPIs from the IID, and considered only those PPIs that (a) involve genes that each have a one-to-one orthologue to the chicken with an orthology confidence score of 1 (using data from Ensembl Compara [139], a score of 1 indicates compliance with the gene tree), a reciprocal % gene identity of >= 75%, a whole genome alignment score of >= 75%, and a gene order conservation score of >= 75% (indicating a high degree of contiguity around the gene of interest), (b) have UniProt IDs that are unambiguously assigned to only one human gene ID (and thereby only one orthologous chicken gene ID), and (c) have PPI evidence type ‘exp’ or ‘pred’.

### Availability of datasets

To test whether down-sampling quantitatively alters the expression profile of an RNA-seq dataset, we randomly down-sampled each of the 18 BMDM datasets (+/- LPS) to 10 million reads 100 times, using seqtk seeded with a random integer between 0 and 10,000. These sets of expression estimates are available as Dataset S1, hosted on the University of Edinburgh DataShare portal (http://dx.doi.org/10.7488/ds/2137). The meta-atlas of chicken gene expression is available in full as Table S6 and via the cross-species annotation portal BioGPS (http://biogps.org/dataset/BDS_00031/chicken-atlas/). To compare genes between species and to visualise expression profiles, BioGPS requires that each gene have an Entrez ID, although this is not the case for all genes in GalGal5. The expression profiles of those genes without Entrez IDs can be found in Table S6.

### Analysis of chicken developmental samples

The expression data derived from CAGE [53] were obtained from http://fantom.gsc.riken.jp/5/suppl/Lizio_et_al_2017/data; the expression file is named galGal5.cage_peak_tpm.osc.txt.gz and the annotation file galGal5.cage_peak_ann.txt. The annotation and expression files were emerged based on chromosomal location of the promoter. All promoters where no sample exceeded 10 tags per million (tagsPM) were excluded from the analysis. The expression data were then entered into Miru (as described above), using a correlation coefficient threshold of 0.75. 22,839 nodes joined by 5,035,102 edges were entered into the analysis and clustered with an MCL inflation value of 2.2, resulting in 132 clusters of at least 10 nodes.

## DECLARATIONS

### Availability of data and materials

The datasets generated during this study are available in the European Nucleotide Archive under accessions PRJEB22373 and PRJEB22580. All data analysed during this study are included in this published article (and its supplementary information files). The atlas of chicken gene expression is also available via the cross-species annotation portal BioGPS (http://biogps.org/dataset/BDS_00031/chicken-atlas/).

## Authors’ contributions

DAH coordinated the study. LF, AJM, and JOD performed macrophage cell culture and RNA extraction. AP, JS and MS, funded, generated and provided RNA-seq data from the caecal tonsils of *Campylobacter*-infected birds. CW and CA prepared data for visualisation with BioGPS. SJB performed all bioinformatic analyses with the exception of the CAGE analysis. KMS performed the CAGE analysis. SJB and DAH wrote the manuscript. All authors read, contributed to, and approved the final manuscript.

### Consent for publication

Not applicable.

### Ethics approval and consent to participate

Approval was obtained from The Roslin Institute’s and the University of Edinburgh’s Protocols and Ethics Committees. All animal work was carried out under the regulations of the Animals (Scientific Procedures) Act 1986.

## SUPPLEMENTAL MATERIAL

Dataset S1. Expression level estimates generated after randomly down-sampling the BMDM (+/- LPS) datasets to 10 million reads 100 times.

Table S1. Data sources for creating an RNA-seq meta-atlas.

Table S2. Independent datasets sequencing the same tissue/cell type.

Table S3. Exponents of the log-log plots after plotting the reverse cumulative distribution of TPM per gene on a log-log scale.

Table S4. Number of genes with detectable expression, per tissue, after the first iteration of Kallisto.

Table S5. Transcripts not detectably expressed (at > 1 TPM) in any tissue, after the first iteration of Kallisto.

Table S6. Chicken RNA-seq meta-dataset, after the second (and final) iteration of Kallisto.

Table S7. Proportion of RNA-seq reads retained by down-sampling the LPS-stimulated BMDM datasets.

Table S8. Number of detectably expressed genes after randomly down-sampling the LPS-stimulated BMDM datasets.

Table S9. Range of expression estimates, and absolute difference between largest and smallest estimate, after randomly down-sampling the LPS-stimulated BMDM datasets.

Table S10. GO term enrichment for those subsets of genes whose highest PEM is for a given tissue.

Table S11. All-against-all correlation matrix for each tissue in the meta-dataset.

Table S12. Tissues whose expression vectors are most strongly correlated with each other.

Table S13. Clusters of co-expressed genes (obtained via network analysis of the RNA-seq meta-dataset), including candidate gene names for unannotated GalGal5 protein-coding genes.

Table S14. Proportion of genes in each co-expression cluster whose highest PEM is for a given tissue.

Table S15. GO term enrichment for co-expression clusters containing >= 100 genes.

Table S16. Correlation of expression profiles for genes with a known protein-protein interaction.

Table S17. Clusters of co-expressed CAGE tags, obtained via network analysis of the Lizio, *et al* dataset [53].

Table S18. Comparison of co-expression clusters between the RNA-seq atlas and the Lizio, *et al* CAGE dataset [53].

